# Emerin loss of function inhibits MyoD-driven differentiation of human iPSCs into skeletal myotubes

**DOI:** 10.64898/2026.09.14.750687

**Authors:** Tracy A. Knight, Jessica Mella, Daniel Chen, Rodolfo S. Ferreira, Helen C. Miranda, Abigail Buchwalter

## Abstract

Loss of function of the nuclear lamina-associated protein emerin causes Emery-Dreifuss muscular dystrophy (EDMD). Efforts to define emerin’s essential functions in skeletal muscle have been limited by poor concordance between mouse models and human disease phenotypes. Here, we adapt transgene-driven differentiation of human induced pluripotent stem cells (hiPSCs) into skeletal muscle (iSMs) as a tractable human model for emerin loss of function. We find that *EMD* knockout (KO) hiPSCs are poorly responsive to combined overexpression of MyoD and Baf60c and produce fewer mature iSMs, indicating that emerin influences muscle differentiation downstream of these differentiation factors. While MyoD acetylation and heterodimerization with E-box proteins are unaffected by emerin loss, MyoD targets including p21 and myogenin are downregulated, and *EMD* KO cells exhibit impaired cell cycle exit in response to differentiation signals. Transcriptomic analysis of *EMD* KO iSMs revealed persistent expression of cell cycle genes and decreased expression of terminal muscle differentiation genes. Dysregulated genes do not overlap with lamina-associated domains (LADs) but are instead enriched for targets of polycomb repressive complex 2 (PRC2), which deposits H3K27 trimethylation. Altogether, our data indicate a functional overlap between emerin and PRC2-mediated regulation of terminal muscle differentiation.

## Introduction

Emery-Dreifuss muscular dystrophy (EDMD) is the third most common hereditary muscular dystrophy, and is characterized by muscle wasting, limb contractures, and cardiomyopathy.^1^ EDMD is caused by mutations to several proteins that make up a nuclear structure known as the nuclear lamina. The nuclear lamina protects the nucleus from external forces, connects the nucleus to the cytoskeleton, and scaffolds chromatin organization.^2^ The lamina is composed of the lamins, polymer-forming intermediate filament proteins that form a meshwork underneath the inner nuclear membrane (INM). The lamin meshwork is functionalized and bound by various soluble and membrane proteins. While the nuclear lamina exists in some form in all mammalian cell types, cardiac and skeletal muscle are uniquely vulnerable to mutations to lamina components. Mutations to the *LMNA* gene or to genes that encode lamin-binding proteins (*EMD, SYNE1*) cause EDMD and/or other related syndromes.^3^

Emerin is a broadly expressed inner nuclear membrane (INM) protein and an important component of the nuclear lamina. Loss-of-function mutations to the *EMD* gene, which is located on the X chromosome, cause an X-linked recessive form of EDMD.^1,4^ Emerin is a member of a small family of INM proteins that share a conserved LAP2β-emerin-MAN1 (LEM) domain fold that binds to the DNA-crosslinking protein BAF (barrier-to-autointegration factor; also known as BANF1).^5^ Emerin also binds directly to lamin A/C via an unstructured domain to form a ternary emerin-BAF-lamin A/C complex.^6^ In addition to participating in this high-affinity complex, emerin has been linked to many other cytoskeletal, signaling, and chromatin components.^7–10^

It has proven difficult to place emerin’s functions into the disease-relevant context of skeletal muscle. Attempts to define the consequences of emerin loss of function in murine models have delivered mixed results. The *Emd^-/y^* mouse does not develop muscular dystrophy and instead exhibits only subtle cardiac and motor coordination phenotypes,^11,12^ yet *Emd^-/-^* muscle progenitors exhibit differentiation defects when cultured *ex vivo*.^12–14^ These inconsistencies may be due to functional compensation by lamina-associated polypeptide 1 (LAP1), which is more highly expressed than emerin in murine, but not human, muscle.^15^ To understand the unique vulnerability of human skeletal muscle cells to emerin loss of function, we require a tractable human model system.

Embryonic development of skeletal muscle begins with mesodermal specification, followed by the development of myogenic progenitors.^16^ The transcription factors MyoD and Myf5 drive commitment of progenitors to myoblasts and induce the myogenin transcription factor, which in turn drives terminal differentiation of myoblasts into multinucleated myotubes.^16^ Post-natal muscle growth and repair both involve the activation of a tissue-resident pool of quiescent muscle progenitors, often referred to as satellite cells. Upon activation, satellite cells express MyoD and briefly re-enter the cell cycle before terminally differentiating in response to myogenin expression.^16^ MyoD and Myf5 have overlapping functions in myoblast fate commitment,^17^ while myogenin is uniquely essential for terminal muscle differentiation.^18,19^ This cell cycle to terminal differentiation transition is critically mediated by Polycomb Repressive Complex 2 (PRC2). PRC2 methylates histone 3 at lysine 27 (H3K27), which promotes heterochromatin compaction and decreases target gene expression. It is necessary to prevent premature differentiation in cycling cells as well as inhibit cell cycle re-entry in differentiated myotubes.^20–22^ BAF60C, a subunit of the SWI/SNF chromatin remodeling complex, is highly expressed in skeletal muscle and additionally plays a critical role in facilitating MyoD-dependent gene transcription. BAF60C is also notably not present in pluripotent cells.^23–25^

Emerin has been implicated in the regulation of MyoD and Myf5 expression,^26^ suggestive of a role at the myoblast commitment stage. On the other hand, cultured *Emd^-/y^* mouse myoblasts exhibit impaired terminal differentiation propensity,^12–14^ suggesting a role in the terminal differentiation cascade downstream of earlier MyoD and Myf5 function. Here, we directly test emerin’s role in terminal muscle differentiation by adapting a transgene-driven directed differentiation protocol in human induced pluripotent stem cells (hiPSCs).^25,27^ In this induced skeletal muscle (iSM) approach, ectopic expression of MyoD and BAF60C in hiPSCs bypasses mesodermal differentiation and induces direct commitment to the myogenic lineage and formation of terminally differentiated, multinucleated myotubes. We generate iSMs from *EMD^-/y^* and *LMNA^-/-^*hiPSCs and demonstrate that loss of either *EMD* or *LMNA* blunts the ability of MyoD to drive myotube differentiation.

## Results

### Establishing a human induced skeletal muscle (iSM) culture model

Directed differentiation of hiPSCs is a powerful approach for modeling the functions of disease-linked genes in relevant cell types.^28^ In this study, we establish a tractable *in vitro* model of emerin loss of function in human muscle. As cultured *Emd^-/y^* mouse myoblasts exhibit impaired terminal differentiation propensity,^12–14^ we hypothesized that loss of emerin impairs the progression of human myoblasts through the terminal differentiation cascade downstream of MyoD and Myf5. To test this prediction, we adapted a transgene-driven induced skeletal muscle (iSM) protocol in human induced pluripotent stem cells (hiPSCs).^25,27^ In this approach, ectopic expression of MyoD and the BAF60C subunit of the SWI/SNF chromatin remodeler in hiPSCs bypasses mesodermal differentiation and induces direct commitment to the myogenic lineage and subsequent formation of terminally differentiated, multinucleated myotubes (Fig. 1A). We constructed a single tetracycline-inducible cassette containing transgenes encoding the mouse MyoD and human BAF60C2 open reading frames separated by a P2A sequence to ensure proportional expression of each iSM-driving transgene. This cassette was then randomly integrated into the genome of WTC-11 hiPSCs by PiggyBac-mediated transposition.

**Figure 1.**
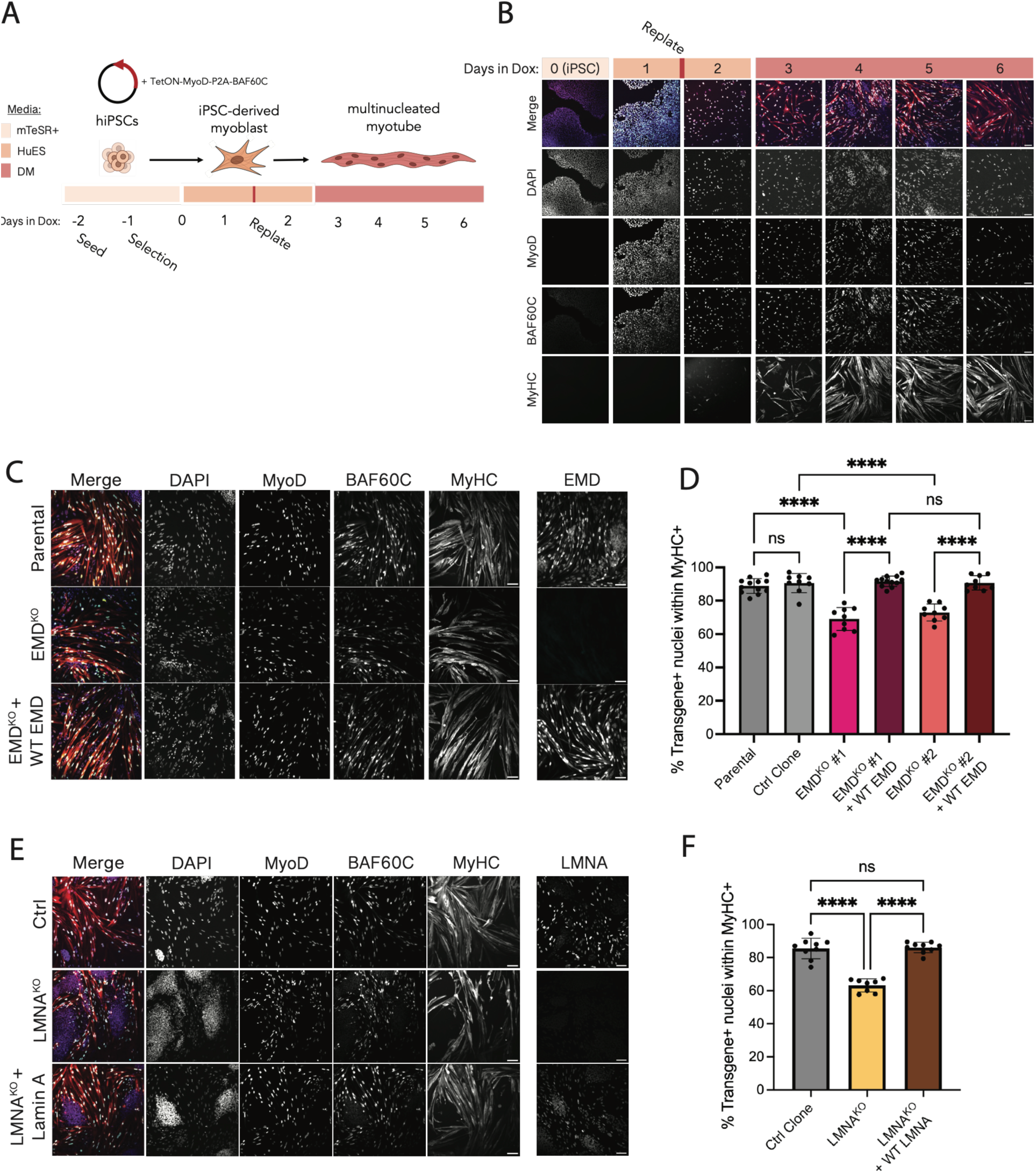
Emerin loss impairs iSM differentiation. A) Diagram of the iSM differentiation protocol. B) Time course of WT (parental) DICE iSM differentiation, following the protocol described in (A). C) Representative images of Day 6 iSMs upon emerin KO and rescue with untagged emerin. D) Quantification of transgene-positive nuclei within MyHC-positive myotubes in (C) (n = 3 biological replicates per genotype). E) Representative images of Day 6 iSMs upon LMNA KO and rescue with untagged Lamin A. F) Quantification of transgene-positive nuclei within MyHC-positive myotubes in (E) (n = 3 biological replicates per genotype). All scale bars = 100um. Proportions significantly different by one-way ANOVA: ****p < 0.0001.

To induce myotube formation, cells were grown in HuES medium with doxycycline for 2 days, then replated and grown in differentiation medium with doxycycline for up to 5 days (see Methods) (Fig. 1A). Earlier versions of iSM protocols involved a 2-day pulse of doxycycline in hESCs followed by embryoid body aggregation.^25^ In hiPSCs, consistent with more recent reports by the protocol developers^27^ and others,^29^ we find that efficient iSM induction requires constitutive induction of MyoD and BAF60C expression throughout the time course. Doxycycline treatment induces the appearance of myosin heavy chain (MyHC)-expressing cells within 3 days that fuse and mature into multinucleated, elongated myotubes over the next several days (Fig. 1B). After 6 days of iSM induction, approximately 90% of transgene-positive nuclei reside within MyHC-positive myotubes (Fig. 1D), demonstrating efficient induction of skeletal muscle differentiation.

Proliferating clusters of transgene-negative cells were sometimes observed (for example, see DAPI-stained, transgene-negative clusters at day 4 and day 5, Fig. 1B). As these cells lack the MyoD/BAF60C transgene, they presumably remain in a stem-like state and continue to proliferate. These cells can be identified as transgene-negative by MyoD/BAF60C immunostaining and microscopy and excluded from further analyses. We determined that the pluripotency-specific cell surface marker CD57^30^ is highly expressed by transgene-negative stem cell clusters, but not by transgene-positive mononucleated cells or mature myotubes (Suppl. Fig. 1) and confirmed that CD57 immunodepletion specifically removed stem cells from differentiated populations, yielding enriched populations of differentiated iSMs (Suppl. Fig. 1) This approach enables bulk analyses without contamination by transgene-negative cells.

### Establishing EMD-null and EMD rescue hiPSCs

EDMD1 exhibits recessive inheritance,^1^ and numerous truncating, frameshift, or missense mutations throughout the *EMD* gene are anticipated to destabilize the *EMD* transcript and/or protein.^4,31^ These features are characteristic of a loss-of-function disease mechanism. We therefore reasoned that *EMD* knockout (KO) would be an effective means to model EDMD1 muscular phenotypes *in vitro*. We also desired a means to efficiently conduct rescue experiments by re-introducing *EMD* or other genes of interest into *EMD* KOs. We accomplished both goals by CRISPR/Cas9 editing as described previously^32^ and summarized here. First, we introduced a dual integrase cassette exchange (DICE) landing pad^33^ into one allele of the *AAVS1* safe harbor locus^32,34^ to generate WTC-11 DICE landing pad hiPSCs (referred to as parental). Next, we knocked out the *EMD* locus in the DICE hiPSCs, selecting two validated *EMD* KO DICE clones for further analysis (Suppl. Fig. 2). We also selected an *EMD* WT CRISPR control clone for comparison (Suppl. Fig. 2). Finally, we confirmed that insertion of the emerin ORF into the *EMD* KO DICE landing pad resulted in constitutive rescue of emerin to a level slightly exceeding endogenous expression (Suppl. Fig. 2).

### EMD-null and LMNA-null hiPSCs are less responsive to skeletal muscle induction

To evaluate the responsiveness of *EMD* KO cells to MyoD/BAF60C-induced differentiation, we integrated the iSM cassette into each clone of *EMD* KO hiPSCs and induced iSM differentiation with doxycycline. In both *EMD* KO clones, we noted a significant decrease in the proportion of transgene-positive cells that produced MyHC-expressing myotubes compared both to WT parental hiPSCs and to a CRISPR control hiPSC clone (Fig. 1C,D; Suppl Fig. 3); we further validated this phenotype in two independent *EMD* KO clones (Suppl. Fig. 4). This myogenic defect is not a consequence of reduced transgene expression, as MyoD and BAF60C are expressed at comparable levels in both genotypes (Suppl. Fig 3). Further, exogenous (mouse) MyoD efficiently induced endogenous (human) MyoD transcript expression in both WT and *EMD* KO cells (Fig. 4B), indicating that MyoD can drive its own feed-forward activation^35^ in both contexts. Importantly, iSM differentiation of *EMD* KO clones is fully rescued by re-expression of WT emerin via DICE integration at the *AAVS1* locus (Fig. 1C,D; Suppl. Fig 3).

As mutations to both *EMD* and *LMNA* cause EDMD, we next compared the consequences of *LMNA* ablation on iSM differentiation. To achieve this, we generated *LMNA* KO DICE landing pad hiPSCs as described above (Suppl. Fig. 2), then integrated the iSM cassette and induced differentiation. Transgene-expressing *LMNA* KOs are similarly impaired in their ability to produce MyHC-expressing myotubes, and this phenotype is again rescued by re-expression of WT Lamin A (Fig. 1E,F). Altogether, these data indicate that *EMD* and *LMNA* each enhance the efficiency of terminal skeletal muscle differentiation induced by MyoD and BAF60C.

### EMD-null iSMs exhibit delayed cell cycle exit

As cell cycle exit normally precedes myotube fusion and MyHC expression, we evaluated proliferation in EMD KO iSMs by pulse-labeling cultures with 5’ethynyl-2’-deoxyuridine (EdU) at each day of the iSM differentiation time course (see Methods) (Fig. 2A). This analysis revealed a two-stage response in WT and EMD rescue iSMs. After replating in differentiation medium, MyoD and BAF60C-expressing cells undergo a drop in their proliferation rate (Fig. 2B-C, compare day 1 to day 2.5; from ∼100% to ∼40% MyoD/BAF60C cells EdU-positive). Over the following days, EdU labeling of transgene-expressing cells continues to decline (∼20% MyoD/BAF60C cells EdU-positive at day 6). In the EMD KOs, the transgene-expressing cells respond similarly to WTs in the first phase (Fig. 2B-C, compare day 1 to day 2.5). However, transgene-expressing *EMD* KOs do not undergo the second phase of cell cycle exit and instead maintain a steady proportion of EdU-positive cells over the remainder of the iSM time course (Fig 2B-C; ∼40% MyoD/BAF60C cells remain EdU positive through day 6). This abnormal proliferation was observed only in MyHC-negative cells, while MyHC-positive myotubes were nonproliferative (Fig. 2B, inset).

**Figure 2.**
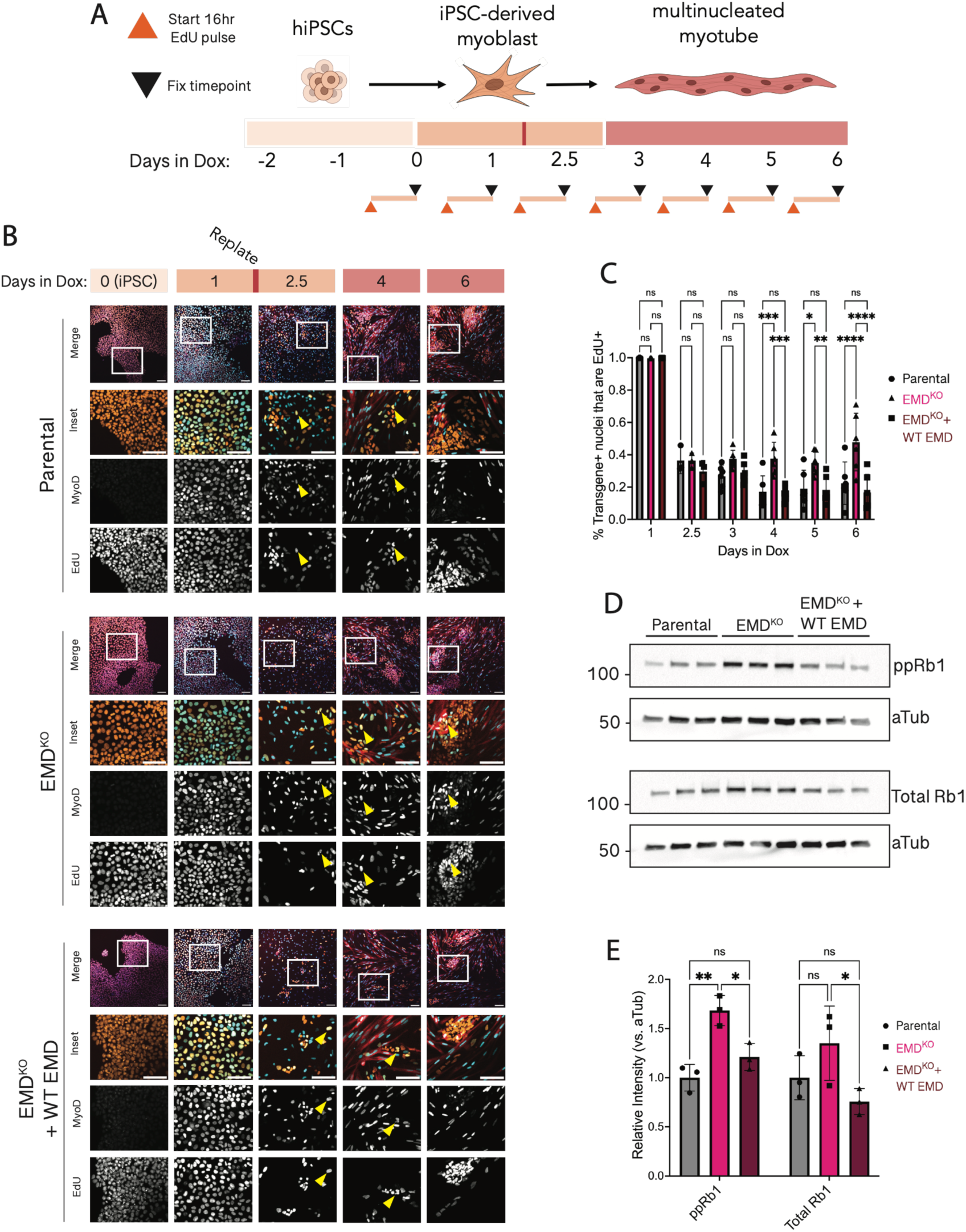
Transgene-positive nuclei exhibit delayed cell cycle exit upon emerin loss. A) Diagram of the EdU labeling protocol. B) Time course of parental wildtype, EMD KO, and EMD KO rescued with untagged WT EMD co-stained for MyoD, MyHC, and incorporated EdU after 16hr pulse. Yellow arrowheads indicate transgene+/EdU+ population, quantified in (C). C) Quantification of transgene-positive nuclei that are also EdU-positive (n = 2 biological replicates per genotype). D) Western blot of hyper-phosphorylated Rb1 (ppRb1) and total Rb1 in Day 6 iSMs. Three biological replicates per genotype. E) Densitometric quantification of (D) as compared to acetylated Tubulin loading control. All scale bars = 100um. Proportions significantly different by two-way ANOVA: *p < 0.05, **p < 0.01, *** p < 0.001.

These results are consistent with prior reports in the *Emd^-/y^*mouse model, where regenerating *Emd^-/y^* muscle and cultured *Emd^-/y^* myoblasts each showed transcriptional signatures of impaired cell cycle exit, including elevated expression of E2F target genes.^12^ E2F-family transcription factors are normally inactivated by interaction with hypo-phosphorylated Rb during cell cycle exit. *Emd^-/y^* mouse myoblasts exhibit elevated levels of hyper-phosphorylated Rb,^12^ implying that Rb-mediated cell cycle exit may be impaired by emerin loss. To test whether Rb is also dysregulated in *EMD* KO human iSMs, we analyzed enriched day 6 iSMs. These analyses revealed a moderate accumulation of hyperphosphorylated Rb in *EMD* KO iSMs that was reverted in *EMD* rescue iSMs (Fig. 2D-E).

MyoD promotes cell cycle exit by driving expression of the Cdk-inhibitory protein p21 (encoded by the *CDKN1A* locus).^36,37^ We tracked p21 induction over each day of differentiation in WT, *EMD* KO, and *EMD* rescue iSMs (Fig. 3A). Replating in differentiation medium quickly induced p21 expression in MyoD/BAF60C transgene-expressing cells, but to a lesser extent in *EMD* KOs than in WT or EMD rescue iSMs (Fig. 3A-B, compare day 1 to day 2). Levels of p21 expression remained persistently lower in *EMD* KOs through the differentiation time course (Fig. 3A-B, days 4 through 6). Similarly, transgene+ cells expressed less p21 in EMD KOs than WT over the course of differentiation (Fig. 3C). This effect was also apparent in bulk measurements of p21 expression in enriched iSM populations at both the RNA (Fig. 3D) and, more variably, at the protein level (Fig. 3E; Suppl. Fig. 3). This blunted p21 expression mirrors the persistence of EdU-labeled proliferating cells in *EMD* KO iSMs (Fig. 2). We also find that the expression of p57 (encoded by the *CDKN1C* locus), a related CIP that functions along with p21 in muscle differentiation, is moderately decreased.^38^ These findings imply that *EMD* KO iSMs are unable to establish and/or maintain cell cycle exit in response to pro-differentiation signals.

**Figure 3.**
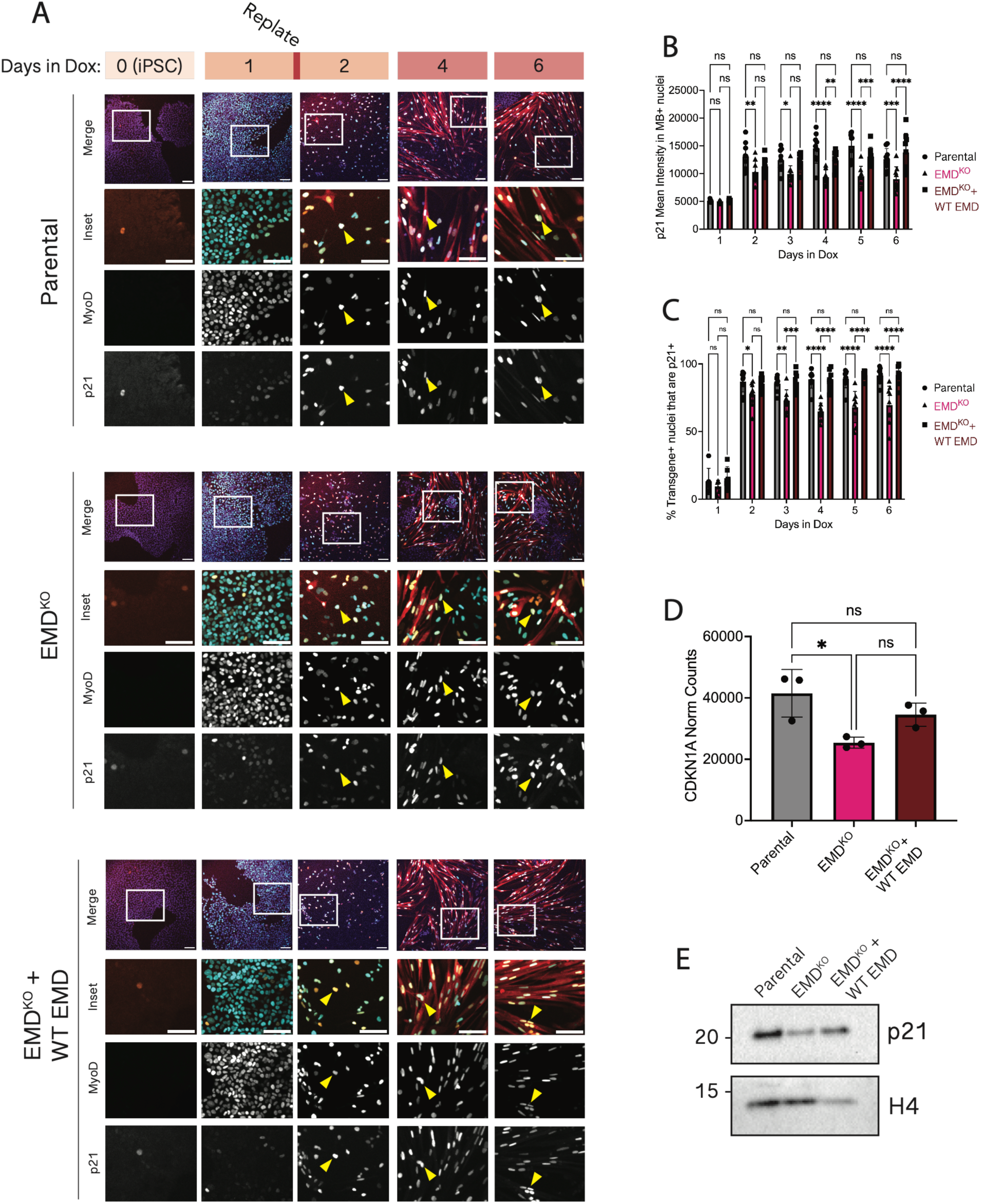
p21 expression is diminished in iSMs upon emerin loss. A) Time course of iSM differentiation, co-stained for MyoD, MyHC, and p21. Yellow arrowheads indicate transgene+/p21+ population, quantified in (B). B) Quantification of transgene-positive nuclei that are also p21-positive (n = 3 biological replicates per genotype). C) Quantification of average p21 signal in transgene-positive nuclei (n = 3 biological replicates per genotype). D) Normalized transcript counts of *CDKN1A* in Day 6 iSMs. E) Western blot for p21 in Day 6 iSMs. All scale bars = 100um. Proportions significantly different by two-way ANOVA in (B) and (C), one-way ANOVA in (E): *p < 0.05, **p < 0.01, *** p < 0.001.

### MyoD activation is unimpaired in EMD KO iSMs

The decrease of p21 expression in the absence of emerin (Fig. 3) could reflect impaired transactivation by MyoD, as the p21-encoding locus *CDKN1A* is a direct transcriptional target of this TF.^36,37,39^ MyoD’s activity is controlled by a post-translational modification switch: Cdk phosphorylation in proliferating cells restrains MyoD activity, while acetylation promotes MyoD activity.^40,41^ MyoD acetylation can accumulate only in the absence of inhibitory phosphorylation.^41^ To explore emerin’s potential influence on MyoD regulation, we immunoprecipitated MyoD from day 6 iSMs. Surprisingly, MyoD was robustly acetylated in WT, *EMD* KO, and *EMD* rescue iSMs (Fig. 4A). MyoD also forms a high-affinity, transcriptionally active heterodimer with the E12/E47 (also known as E2A) E-box proteins^42^; we detect this robust interaction equivalently in WT, *EMD* KO, and *EMD* rescue iSMs (Fig. 4A). These data, along with the observation that the (mouse) MyoD transgene can effectively transactivate the endogenous (human) MyoD locus in each genotype (Fig. 4B), imply that emerin loss does not impair MyoD activation.

**Figure 4.**
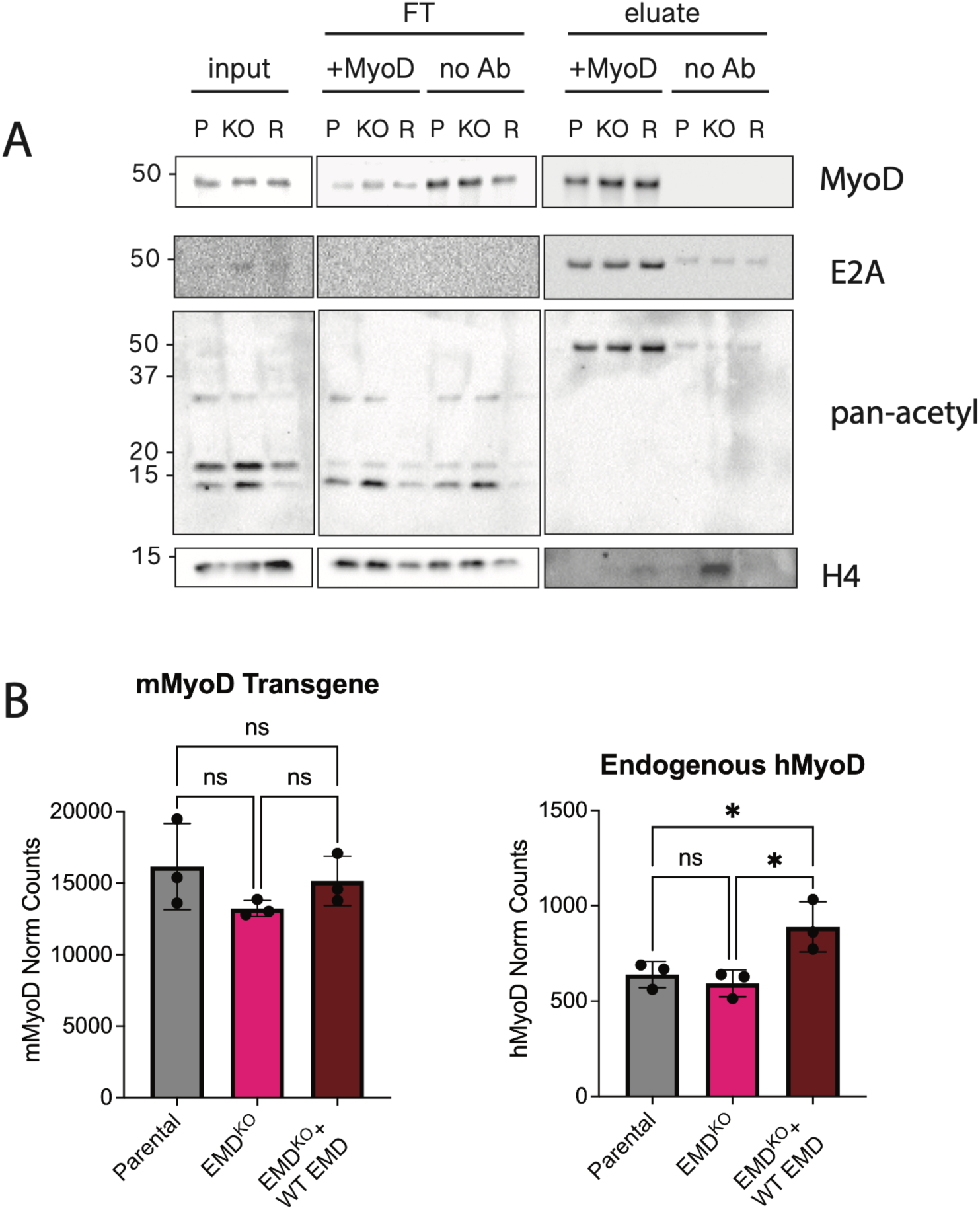
MyoD activation is unaffected by emerin loss. A) Representative immunoblots for immunoprecipitation of MyoD and co-immunoprecipitation of E2A (repeated with 3 biological replicates). FT = flow-through, P = parental, KO = EMD^KO^ #1, R = EMD^KO^ #1 + WT EMD (Rescue). B) Normalized transcript counts of the mouse *Myod1* transgene and endogenous human *MYOD1*. Counts significantly different by one-way ANOVA: *p < 0.05.

### Myogenin expression is decreased in EMD KO iSMs

Myogenin is a late, direct transcriptional target of MyoD^18,19^ that is essential for terminal muscle differentiation.^16^ We noted that *MYOG* transcript levels were decreased in *EMD* KO iSMs and rescued by re-expression of *EMD* (Suppl. Fig. 5D). We therefore evaluated myogenin expression over the course of differentiation in WT, *EMD* KO, and *EMD* rescue iSMs (Suppl. Fig. 5A). Transgene+ nuclei did not fully co-express myogenin until the final days of differentiation, consistent with the role of this transcription factor in terminal differentiation. Transgene+ nuclei expressed myogenin in similar proportions in WT, *EMD* KO, and *EMD* rescue iSMs (Suppl. Fig. 5B). However, there was a small but significant decrease in the proportion of myogenin+ nuclei contributing to MyHC+ myotubes in *EMD* KOs (Suppl. Fig. 5C), comparable to the proportional decrease of transgene+, MyHC+ nuclei that we observed in *EMD* KOs (Fig. 1). These results indicate that *EMD* loss moderately impairs the expression and/or activity of myogenin.

### Transcriptional signatures of impaired cell cycle exit and differentiation in EMD KO iSMs

To probe the effects of *EMD* loss on the iSM transcriptome, we performed RNA-sequencing (RNAseq) on three biological replicates of enriched day 6 WT, *EMD* KO, and *EMD* rescue iSMs. There was a strong, positive correlation (R^2^ = 0.737) between genes differentially expressed in *EMD* KO iSMs and genes responsive to *EMD* rescue, indicating that re-introduction of *EMD* generally brings expression of these genes back towards a WT state (Fig. 5A). In total, we identified 626 DEGs that were differentially expressed by at least 1.5-fold in *EMD* KOs and rescued by re-introduction of *EMD*, which we refer to hereafter as rescue-responsive genes. Gene Set Enrichment Analysis (GSEA) revealed enrichment of cell cycle regulators among genes upregulated in *EMD* KOs (Fig. 5B), such as *AURKA, CCNA2,* and *CDK1,* consistent with our observations of impaired cell cycle exit (Fig. 2). We also observe upregulation of homeobox genes critical to embryonic development like *DLX4* and *SIX3* (Fig. 5A). Amongst genes downregulated in *EMD* KOs, GSEA identified factors related to skeletal muscle function and terminal differentiation (Fig. 5B), including muscle-specific myosins *(MYH3, MYH7, MYH7B, MYH8),* transcription factors (*MYOG)*, and CIPs (*CDKN1A, CDKN1C)* (Suppl. Fig. 6).

**Figure 5.**
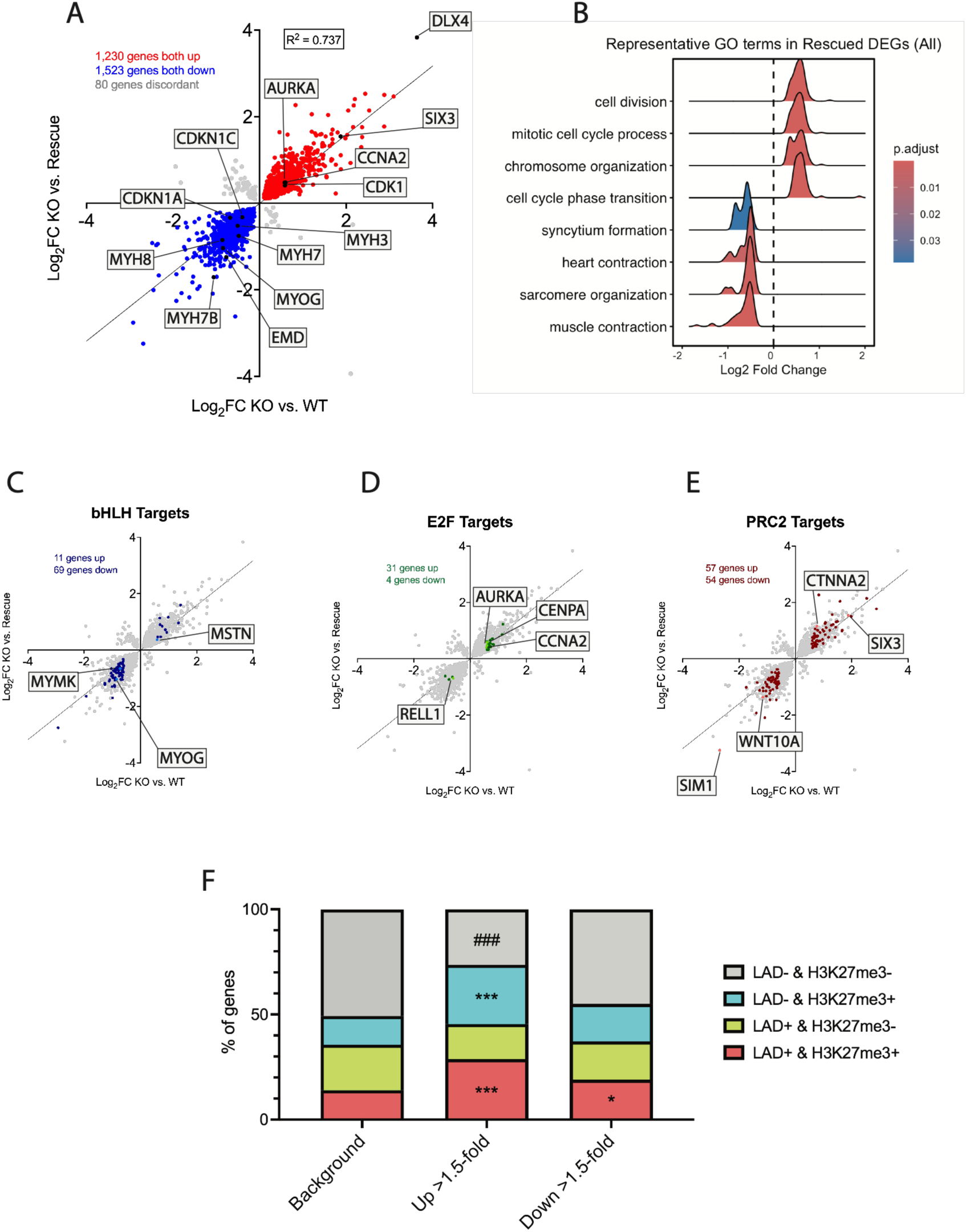
Emerin loss leads to upregulation of proliferative genes and downregulation of myogenic differentiation genes. A) Identification of emerin-sensitive genes. Log_2_ fold-change (FC) values in EMD^KO^ vs. Parental (KO vs. WT) plotted against log_2_FC values in EMD^KO^ vs. EMD^KO^ + WT EMD (KO vs. Rescue). Best-fit shown as solid black line (y = 0.7939x – 0.005017; R^2^ = 0.737). Genes of interest labeled in black with gene names. B) Representative GO terms enriched among all emerin-sensitive genes shown in (A) (identified by gene set enrichment analysis). C-E) Identification of enriched transcription factor (TF) targets among emerin-sensitive genes. Genes identified as targets of the bHLH TF family (C), the E2F-DREAM TF family (D), and the PRC2 complex (E) by the ChIP-Atlas database are highlighted against the same scatterplot shown in (A). Representative target genes are labeled. Also see Supplementary Fig. 7. F) Distribution of rescue-responsive genes among LAD and/or H3K27me3 regions reported in human myotubes. Background corresponds to 20,036 protein-coding genes found in both published datasets. Proportions significantly different from background by two-sided Fisher’s exact test: *p < 0.05, ***p < 0.001 (increased) and ###p < 0.001 (decreased).

### Dysregulated genes are not enriched in lamina-associated domains (LADs)

The nuclear lamina scaffolds heterochromatic lamina-associated domains (LADs) which cover approximately 40% of the genome. Due to emerin’s association with the lamina, we hypothesized that rescued genes may be enriched within LADs. To assess this, we compared our list of rescue-responsive genes to Lamin A and Lamin B1 genome-binding datasets deposited in the ChIP-Atlas database.^43^ Surprisingly, this analysis did not indicate an enrichment of rescued genes within LADs associated with either Lamin A or Lamin B1.

While many LADs are constitutive, their genomic coverage does vary across cell types. Therefore, to confirm this result in muscle cells, we compared our list of rescue-responsive genes with published DamID LAD maps from murine C2C12 myoblasts and myotubes^44^ and from human myotubes and myo-converted fibroblasts.^45^ These comparisons indicated that upregulated rescue-responsive genes were modestly enriched within LADs (Suppl. Fig. 7). However, as many dysregulated genes are not found within LADs, we conclude that gene expression changes induced in *EMD* KO iSMs cannot be explained solely by LAD disruption.

### E2F, bHLH, and PRC2 targets are enriched among dysregulated genes in EMD KO iSMs

To conduct an unbiased search for candidate transcription factors or co-regulators that underlie differential gene expression in *EMD* KO iSMs, we compared our list of rescued genes to the full breadth of genome-binding datasets in the ChIP-Atlas database: 1,597 targets in total, including various transcription factors, chromatin modifiers, and chromatin architectural proteins. Top factor categories targeting these genes included the bHLH family of myogenic transcription factors, the E2F-DREAM cell cycle regulatory pathway, and polycomb repressive complex 2 (PRC2) (Fig. 5C-E). bHLH TF targets included genes involved in terminal muscle differentiation, which were mainly downregulated; in contrast, E2F targets included a cluster of upregulated, pro-proliferative genes. PRC2 targets included both up- and down-regulated genes. These findings suggest a general defect in PRC2-dependent gene regulation in *EMD* KO iSMs.

### Upregulated genes are enriched for H3K27me3

The PRC2 complex is responsible for the di- and tri-methylation of H3K27.^22^ To further explore the relationship between *EMD* loss and dysregulation of PRC2 target genes, we cross-referenced our rescue-responsive genes to publicly available ChIP-seq datasets mapping H3K27me3 occupancy in human myotubes.^46–48^ This analysis revealed that genes upregulated in *EMD* KOs are strongly enriched for H3K27me3 in normal myotubes (Fig. 5F). Genes downregulated in *EMD* KOs also show moderate enrichment for H3K27me3 (Fig 5F; Suppl. Fig. 6). Both H3K27me3 and lamin associated domains (LADs) demarcate heterochromatic domains,^49,50^ but their correlation with each other is highly cell-type and context dependent.^51–53^ While we noted that upregulated *EMD-*sensitive genes are modestly enriched in myotube LADs, cross-referencing LAD and H3K27me3 datasets revealed that it is actually the subset of LADs that are H3K27me3-modified in myotubes that show a strong enrichment for upregulated *EMD-*sensitive genes, and a moderate enrichment for downregulated genes (Fig. 5F). We also confirmed that LADs identified from these datasets were significantly more enriched for H3K27me3 in human myotubes non-LADs (Suppl. Fig. 7). Altogether, these results demonstrate that H3K27me3 modification, rather than LAD association, is a general shared feature among genes dysregulated by *EMD* loss.

## Discussion

Elucidating the functional role of emerin in human skeletal muscle differentiation and maintenance is critical for understanding the mechanism and potential treatment routes for EDMD1. However, EDMD1 disease modeling has been limited by the lack of a phenotype in *Emd* KO mice. There have been some improvements in EDMD1 animal models recently; a rat knockout model was reported to develop defects in both cardiac and skeletal muscle, consistent with symptoms observed in human patients.^54^ Despite these advances, the field still lacks an efficient model to study this disease in a human context. Here, we establish iSMs as a tractable human model of EDMD1. This model provides an accessible resource to study emerin function in human cells.

The severity of the *EMD* KO phenotype in iSMs is particularly notable in comparison to other cellular models of muscular dystrophy. For instance, a previous study used a similar transgene expression system to induce Duchenne muscular dystrophy (DMD) patient-derived iPSCs to form myocytes and reported minimal differences in differentiation propensity or morphology between controls and DMD mutant myocytes.^29^ However, DMD myocytes still exhibited pronounced abnormalities in calcium influx and secretion of creatine kinase which was rescued by treatment with an exon-skipping antisense oligonucleotide. These DMD models utilized patient-derived iPSCs harboring disease-causing nonsense or missense mutations. Similarly, EDMD1 mutations are destabilizing and characteristically lead to nearly undetectable protein levels in patients, making *EMD* KO a reasonable strategy to model the disease *in vitro*. The clear differentiation defect of *EMD* KO iSMs emphasizes the necessity of emerin for *in vitro* skeletal muscle development.

Previous studies using cellular models have reported a requirement for emerin function at early stages of myogenesis. Emerin has been reported to be required for positioning gene loci encoding the early myogenic differentiation factors Pax7, Myf5, and MyoD at the nuclear periphery.^26^ Emerin has also been implicated in activation of MyoD and its downstream targets in *EMD* KO mouse muscle, despite the lack of an overt muscular dystrophy phenotype.^12^ Here, in contrast, we report a requirement for emerin downstream of MyoD expression, as MyoD overexpression is unable to rescue the overt phenotype caused by emerin loss in human iSMs. In response to MyoD overexpression, *EMD* KO iSMs show persistent DNA replication in MyoD-expressing cells, retain hyperphosphorylated Rb (Fig. 2), and less effectively induce p21 (Fig. 3).

We also demonstrate the strong association between emerin-sensitive genes and PRC2. Several studies have shown that the timing of PRC2 activity is critical in myogenesis both for silencing myogenesis genes to prevent premature differentiation as well as silencing cell cycle genes in late myogenesis to promote terminal differentiation. In satellite cells, activity of the PRC2 component EZH2 maintains the proliferating satellite cell population by limiting expression of myogenesis genes.^55^ Interestingly, the *CDKN1A* locus, which encodes p21, is repressed by EZH2 in cycling C2C12 myoblasts.^56^ Notably, p21 induction is blunted in *EMD* KO iSMs, suggesting that emerin may promote the timely removal of EZH2-mediated repression at this locus during myogenesis (Fig. 3). In late myogenesis, the PRC2 complex is remodeled by EZH2 degradation and replacement with EZH1, a structurally similar but less active H3K27 methyltransferase.^57–59^ In this current study, we find that a loss of emerin leads to improper upregulation of many H3K27me3-marked genes. Surprisingly, we also find limited association between emerin-sensitive genes and myotube LADs. Due to these findings, we speculate that emerin influences PRC2/EZH2 targeting and/or activity during myogenesis, both within and outside of LADs.

We recognize certain limitations to this current study. While iSMs are a valuable system for studying skeletal muscle differentiation in a human context, they bypass native control of lineage specification by ectopic overexpression of MyoD and BAF60C. Additionally, while we undertook comparative studies leveraging existing genomic datasets, it will be valuable to map LADs and H3K27me3 genomic enrichment specifically in iSMs. We are unable to comment on the direct requirement of PRC2 in iSMs due to this limitation – we simply observe that its targets are enriched among genes dysregulated in *EMD* KOs but cannot say whether emerin loss alters PRC2 localization or activity. In conclusion, our work will enable future studies into the essential functions of emerin in human myogenesis.

## Methods

### Cell Culture

WTC-11 hiPSCs (healthy 30-year-old, XY) were obtained from and verified by the Berkeley Stem Cell Center. hiPSCs were cultured at 37C in 5% CO_2_ under normoxic conditions in either mTeSR+ medium (Stem Cell Technologies 100-0274/100-0275) or 1:1 mTeSR+:HiDef-B8 (Defined Biosciences LSS-204) in vessels coated with 1:15 Geltrex (Gibco A14132-02) without penicillin/streptomycin. Cells were passaged as clumps using ReLeSR (Stem Cell Technologies 100-0483) for routine culture and as single cells using Accutase (Innovative Cell Technologies AT-104) for counting and electroporating. Cells were plated in 10uM ROCK inhibitor Y-27632 (Selleck Chemicals 101763-964), then changed to fresh hiPSC media after 24 hours, with media changed ever 48 hours thereafter.

### Generation of DICE hiPSCs

Genome editing was performed by electroporating 1 × 10^6 cells with 12.5 pmol Cas9-NLS protein and 12.5 pmol sgRNA using a 100-µL Neon electroporation system (1300 V, 30 ms, 1 pulse). For DICE landing pad integration, 2 µg of the HDR donor plasmid containing the landing pad cassette was included in the electroporation mixture. Following transfection, cells were selected with 0.25 µg/mL puromycin for 3 days. Single-cell clones were isolated by limiting dilution in 96-well plates, expanded, and genotyped by PCR using primers spanning the AAVS1 homology arms to identify heterozygous landing pad integration.

Emerin knockout cell lines were generated from a single landing pad clone using two sgRNAs targeting the 5′ and 3′ ends of the coding sequence. Deletion of the targeted locus was confirmed by PCR, and loss of emerin expression was verified by immunoblotting and immunofluorescence microscopy. For site-specific integration of emerin constructs, 1 × 10^6 landing pad cells were electroporated with 2 µg of an attB-containing donor plasmid and 500 ng of a BxbI-P2A-PhiC31 integrase plasmid using the same electroporation conditions. After recovery, BFP-negative, donor-positive cells were isolated by FACS using a SONY SH800 cell sorter. Cell lines were routinely tested for mycoplasma every 3–4 months and were consistently negative.

### Generation of iSM DICE hiPSCs

Tetracycline-inducible iSM transgene vector was amplified by PCR from two separate vectors gifted by the lab of Dr. Lorenzo Puri. Final vector included rtTA-Advanced-T2A-BlastR controlled by a UbC promoter and mMyod1-P2A-hBAF60C-FLAG controlled by a tight TRE promoter, both flanked by SV40 poly(A) sequences. These features were all enclosed between 5’ and 3’ piggyBac integration sites. Full vector sequence is available upon request.

Expected sequence was confirmed by whole plasmid sequencing. The iSM transgene vector was introduced into DICE hiPSCs using a 100 µL Neon electroporator set to 1150V, 20ms, and 2 pulses. Cells were placed under Blasticidin selection (Research Products International B12150-0.1) to select for mMYOD1/hBAF60C-positive cells (transgene+) for at least 7 days at 10ug/mL.

### Differentiation

For differentiation into iSMs, hiPSCs were seeded in hiPSC media and allowed to recover under Blasticidin selection before switching to HuES media containing 500ng/mL doxycycline (Alfa Aesar J67043). HuES media was made by mixing Dulbecco’s modified Eagle medium:F12 (Gibco 11330-032), 20% Knockout Serum Replacement (Thermo Scientific 10828-028), 1% non-essential amino acids (Caisson Labs NAL03-100ML), and 0.2% Beta-mercaptoethanol (Gibco 21985-023). After 24hr, cells were replated in HuES + doxycycline + ROCK inhibitor. After another 24hr, media was switched to DM media + doxycycline. DM media was made by mixing Dulbecco’s modified Eagle medium:F12, 1X insulin-selenium-transferrin-ethanolamine (ITS-X; Gibco 51500-056) and 1X N-2 (Gibco, 17502-001). Cells were maintained in DM media + doxycycline until collection or imaging, replacing with fresh media every 48 hr.

iSM cleanup was performed on freshly collected Day 6 iSMs. Fresh cell pellets were suspended in anti-CD57 (BioLegend 393330) diluted in 0.5% BSA in PBS for 90min, rotating end-over-end at 4C. Sheep anti-mouse M280 Dynabeads (Thermo Scientific 11201D) were calibrated in 0.5% BSA briefly before the cell-antibody mixture was added and set to rotate end-over-end at 4C for another 90min. Beads were removed using a magnetic stand and cell pellets were washed in 0.5% BSA before being flash frozen.

### Immunofluorescence and EdU incorporation staining

8-chamber *μ*-slide (ibidi 80806) were coated with 1:15 Geltrex for 10min at 37C. Cells were seeded at the time of replating during differentiation, then fixed in 4% PFA (Electron Microscopy Sciences 15710) in PBS for 5 minutes at room temperature at various time points. Cells were permeabilized in IF buffer (0.1% Triton-X, 0.02% SDS, 10mg/mL BSA in 1X PBS) for 20min at room temperature. Primary and secondary antibodies diluted in IF buffer were incubated for 2hr and 1hr at room temperature, respectively, with IF buffer washes in between. DNA was stained with Hoechst during secondary antibody incubation. See antibody table for list of antibodies used and working concentrations.

After 16hr EdU incubation, cells were fixed as described above. The Click-iT EdU Cell Proliferation Kit for Imaging (ThermoFisher C10338) was used according to supplier’s instructions to label EdU with Alexa-555 fluorophore. The above immunofluorescence staining protocol immediately followed EdU labeling.

### Co-immunoprecipitation & Immunoblotting

Immunoprecipitation was performed by lysing cell pellets in RIPA buffer (50mM Tris-HCl pH 8.0, 150mM NaCl, 1% NP-40, 0.5% sodium deoxycholate, 0.1% SDS, EDTA-free Protease Inhibitor Cocktail (Roche, 11836170001)). Cell lysates were incubated with the mouse monoclonal anti-MyoD antibody (BD 554130) at 1:120, rotating end-over-end overnight at 4C. Lysates were then added to 50uL Sheep anti-mouse M280 Dynabeads (calibrated in RIPA buffer) and incubated rotating end-over-end overnight at 4C. Beads were separated from lysate on a magnetic column, washed three times in IP wash buffer (0.1% BSA, 2mM EDTA in PBS) and eluted in 2X sample buffer (5X = 10%SDS, 25 mM Na3PO4, 50% glycerol, 0.5 M DTT, .05% Bromophenol blue) by boiling at 95C for 10min.

For detection of lamins and membrane-bound proteins, cells were pelleted and lysed in 8 M urea, 75 mM NaCl, 50 mM Tris pH 8.0 and cOmplete, Mini, EDTA-free Protease Inhibitor Cocktail. For detection of phosphorylated and/or soluble proteins, cell pellets were lysed on ice for 10min in buffer containing 1% NP-40 (Sigma I3021), 150mM NaCl, 50mM Tris-HCl pH7.5, 1mM EDTA, cOmplete, Mini, EDTA-free Protease Inhibitor Cocktail, and 2X PhosSTOP Phosphatase Inhibitor Cocktail (Roche, 4906845001). Lysates were run on 4-20% Mini-PROTEAN TGX precast gels (BioRad 4561096) and transferred to a MeOH-activated PVDF membrane for 16hr at 30mA, 4C or for 1hr at 100V, 4C. Blots were probed with anti-EMD, anti-Rb1, anti-alpha tubulin, anti-p21, anti-H4, mouse monoclonal anti-MyoD, rabbit polyclonal anti-MyoD, anti-TCF3/E2A, anti-pan-acetylated lysine, anti-Lamin A/C, anti-BAF60C, anti-MyHC, anti-Myf5, anti-Lamin A/C, and anti-vinculin primary antibodies diluted in 5% milk TBST (see Table 2.2 for additional antibody information). Secondary antibodies were diluted at 1:5000 in 5% milk TBST. For ppRb1 detection, blots were blocked in 5% milk TBST, incubated in with anti-ppRb1 in 5% BSA, followed by anti-Rabbit HRP in 5% BSA. Blots were developed using high-sensitivity ECL substrate (Azure Biosystems AC2103) and imaged using a BioRad ChemiDoc XRS+.

### RNAseq Analysis

Three iSM replicates of each genotype were differentiated in separate wells of 6-well plates, then collected on Day 6 for cleanup as described above. Snap-frozen, enriched iSM pellets were sent to Signios Bio for RNA extraction, library preparation, and sequencing. Paired-end sequencing was performed (read length of 150 bp, ∼40 million reads per library). Raw sequencing reads were mapped to the GENCODE V49 primary assembly human genome build (GRCh38) using STAR (version 2.7.10a): --runMode alignReads --readFilesCommand zcat --clip3pAdapterSeq CTGTCTCTTATA CTGTCTCTTATA --clip3pAdapterMMp 0.1 0.1 –outFileNamePrefix ${base}_ --outFilterMultimapNmax 1 --outReadsUnmapped None --outSAMtype BAM SortedByCoordinate --twopassMode Basic. Bam files were indexed using SAMtools (version 1.10) and used to generate RPKM normalized coverage files using the deepTools2 function bamCoverage (version 3.4.5). A read counts table was generated for all conditions using the subread featureCounts package (version 1.6.4) and GENCODE V49 primary annotation. Read counts table was used as an input for the Bioconductor DESeq2 package (version 1.40.2) to generate normalized counts and perform differential gene expression analysis.

MyoD ortholog analysis was performed independently from all other RNAseq analysis. The mouse Myod1 transgene CDS was appended as a synthetic, single exon chromosome/transcript to both the genome FASTA and the transcriptome FASTA. A transcriptome-only Salmon pufferfish index was built from the combined transcriptome FASTA (Salmon version 1.10.1) to which transcript reads were mapped: salmon quant --index salmon_index/ --libType ISR -- mates1 ${SAMPLE}_R1.fastq.gz --mates2 ${SAMPLE}_R2.fastq.gz --output salmon_quant/${SAMPLE}/ --threads 1 –validateMappings. A quants’ file was generated for each replicate and used as the input for DESeq2. The tximport package was used to perform transcript-to-gene mapping before generating normalized counts (R version 4.4.x).

GSEA was conducted with the gseGO function from the clusterProfiler R package (version 4.14.3) using the “Biological Processes” subontology. Gene set size range was set to a minimum of 5 and a maximum of 5,000, with a Benjamini-Hochberg method-adjusted p-value cutoff of 0.01 and simple permutation n of 10,000.

### TFEA analysis

Rescue-sensitive genes were compared against targets of all human antigens listed in the ChIP-Atlas database with sufficient data (version 3.0).^43^ Background was calculated using the union of 19,032 genes listed as targets of at least one of all 1,597 antigens tested. TF families (i.e. E2F, bHLH, etc.) were assigned to each gene based on enrichment of that gene as a TF family target (if a gene was a target of more than one family, it was assigned to the family with the highest proportion of TFs targeting that gene).

### LAD and H3K27me3 Analysis

Mouse myoblast and myotube LAD data was acquired directly from Robson et al.^44^ Human orthologs were mapped using the biomaRt package to map MGI to HGNC homology (only 1:1 orthologs were used). Human myotube and myo-converted fibroblast LAD data was acquired directly from Benarroch et al.^45^ and used as listed. Background was calculated using all human protein-coding genes from GENCODE v45.

H3K27me3 ChIP-seq data were acquired from ENCODE dataset ENCSR000ATI (human myotubes, Histone ChIP-seq). Pseudoreplicated peaks were used for gene assignment. GENCODE v46 primary assembly basic annotation GTF for GRCh38 was used to build a protein-coding gene list background. Genes were classified as “H3K27me3-marked” within a dataset if at least one peak within the BED file of that dataset overlapped with that gene’s TSS +/− 5kb promoter window.

### Image Analysis & Computation

Images were acquired on a Nikon Ti microscope equipped with a CSU-X1 spinning disk confocal using a 10X 0.45 numerical aperture air objective or an AxioObserver with Apotome (Zeiss). All z-stacks were acquired at 0.3*μ*m steps. Raw 16bit files were saved as ND2 files using Nikon Elements (version 6.10.2). All image processing and quantification was done with Fiji (ImageJ2 release 2.17.0). Images are shown as single slices within a z-stack. MyoD was used to create masks of nuclei to measure mean intensities. Percent positive within transgene+ nuclei quantification was determined by presence of mean signal above a negative background threshold. Manual quantification of the percentage of nuclei within MyHC+ and desmin+ myotubes was performed blinded to experimental condition. For nuclei area quantification, the DAPI channel was analyzed using the Adjust Threshold function of ImageJ, followed by separation of overlapping nuclei using the Watershed command. Graphing and statistical analysis was done using GraphPad Prism (version 11.0.1).

## Acknowledgements

We would like to acknowledge all members of the Buchwalter laboratory for their advice and support through the project period, especially Solène Hervé and Eric Martin for suggesting a cell surface marker-based negative selection strategy for removing contaminating stem cells from skeletal muscle cultures. We acknowledge the following sources of funding: the NSF Graduate Research Fellowship Program (T.K.); the Chan Zuckerberg Biohub Investigator Program, NIH / NIAMS R01AR084208, and NIH / NIGMS R35GM142897 (A.B.).

## Author Contributions

T.K. designed and conducted experiments, analyzed data, and wrote the manuscript. J.M. generated DICE landing pad and EMD KO hiPSCs. D.C., R.F., and H.M. conducted pilot skeletal muscle differentiation assays, shared reagents, and advised on adapting this method. A.B. analyzed data, oversaw the project, provided funding support, and wrote the manuscript.

---

**Supplementary Figure 1.**
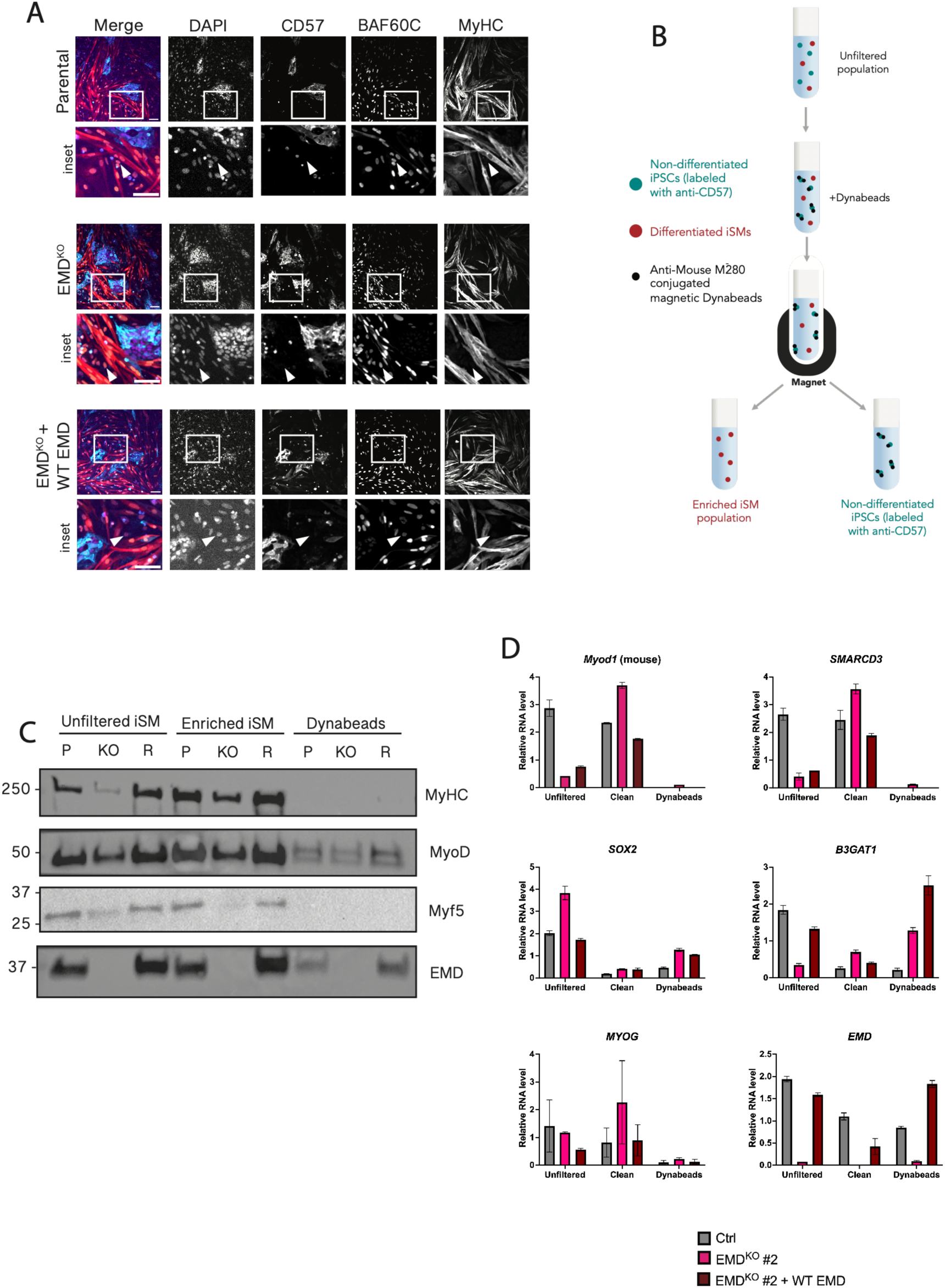
Validation of iSM immunodepletion protocol. A) Immunofluorescence staining of Day 4 iSMs, co-stained for BAF60C, MyHC, and CD57. White arrows indicate examples of BAF60C+/CD57-nuclei. B) Diagram detailing iSM cleanup protocol. C) Western blot of myogenic factors and emerin in unfiltered and enriched iSM populations, as well as CD57+ cells captured by Dynabeads. P = parental, KO = EMD^KO^ #1, R = EMD^KO^ #1 + WT EMD (Rescue). D) RT-qPCR of select genes in CRISPR control clone, EMD^KO^ #2, and EMD^KO^ #2 + WT EMD Day 6 iSMs, comparing unfiltered, enriched, and Dynabead-bound populations (n = 3 technical replicates per condition). RNA level calculated relative to GAPDH control. Gene transcripts quantified: *Myod1* (mouse MyoD transgene), *SMARCD3* (encoding BAF60C), *SOX2*, *B3GAT1* (encoding CD57 synthase), *MYOG* (encoding myogenin), and *EMD*. All scale bars = 100um.

**Supplementary Figure 2.**
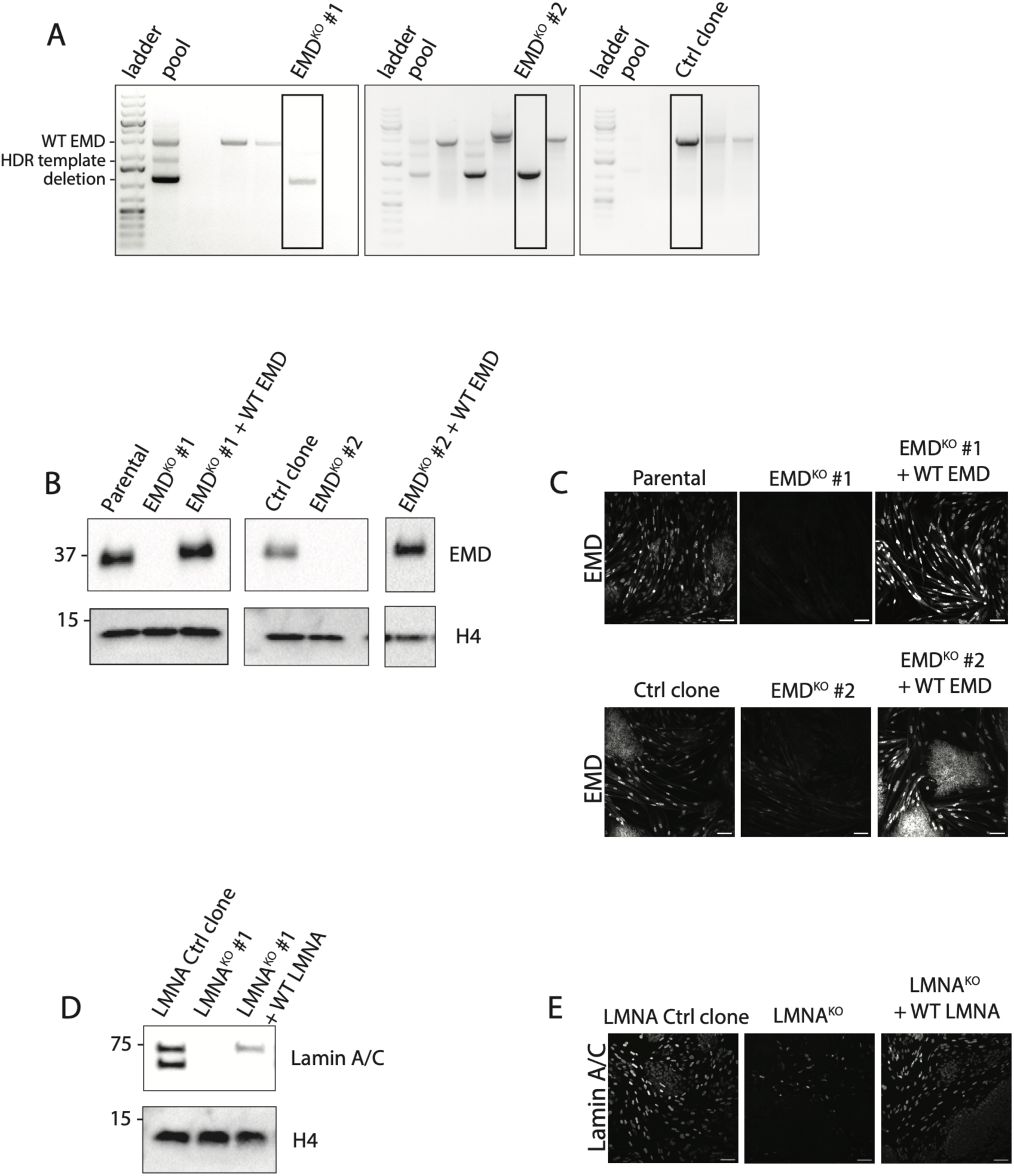
Validation of *EMD* KO and *LMNA* KO hiPSCs. A) Genotyping PCR for *EMD* KO #1, *EMD* KO #2, and CRISPR control. Upper band corresponds to wildtype emerin locus, lower band corresponds to deletion within the *EMD* locus. B) Western blot of CRISPR control clone and both *EMD* KO clones with corresponding rescue. C) Immunofluorescence staining of CRISPR control clone and both *EMD* KO clones with corresponding rescue. D) Western blot of control, *LMNA* KO, and *LMNA* KO + Lamin A rescue, blotted for Lamin A/C. E) Immunofluorescence staining of control, *LMNA* KO, and *LMNA* KO + Lamin A rescue. All scale bars = 100um.

**Supplementary Figure 3.**
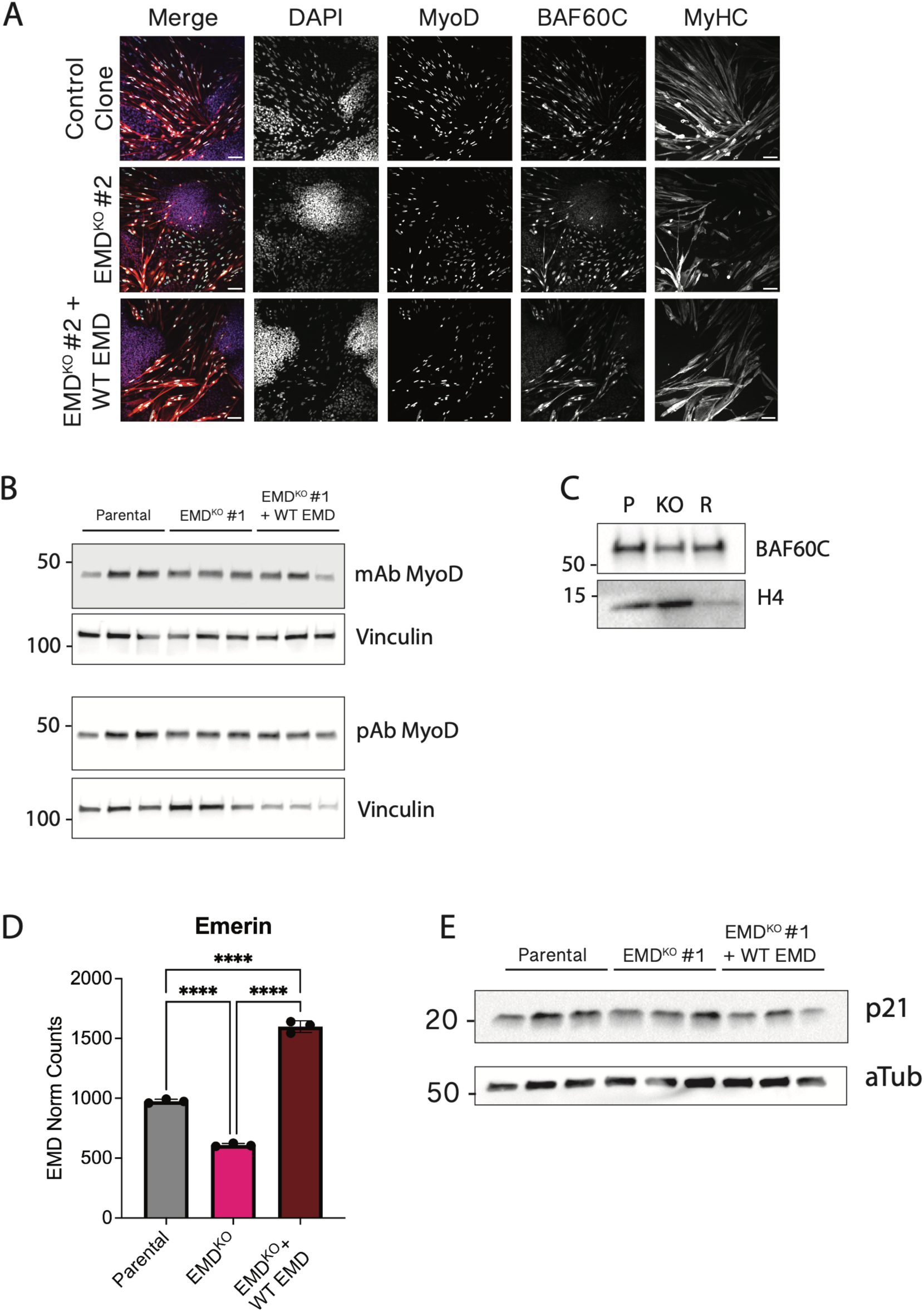
Additional EMD^KO^ iSM differentiation defects and characterization of transgene, *EMD* transcript, and variable p21 levels. A) Representative images of Control, EMD KO #2, and EMD KO #2 + WT EMD Day 6 iSMs, quantified in Figure 1D. B) Three distinct biological replicates immunoblotted using mouse monoclonal (BD cat#) and rabbit polyclonal (Ptg cat#) anti-MyoD antibodies. C) Immunoblot of BAF60C levels. P = parental, KO = EMD^KO^ #1, R = EMD^KO^ #1 + WT EMD (Rescue). D) Normalized transcript counts of *EMD* in Parental, EMD^KO^ #1, and EMD^KO^ #1 Rescue. E) Three distinct biological replicates immunoblotted to demonstrate p21 variability by Western blot (compare with Figure 3E). All samples in this figure are from Day 6 iSMs. All scale bars = 100um. Proportions significantly different by one-way ANOVA: ****p < 0.0001.

**Supplementary Figure 4.**
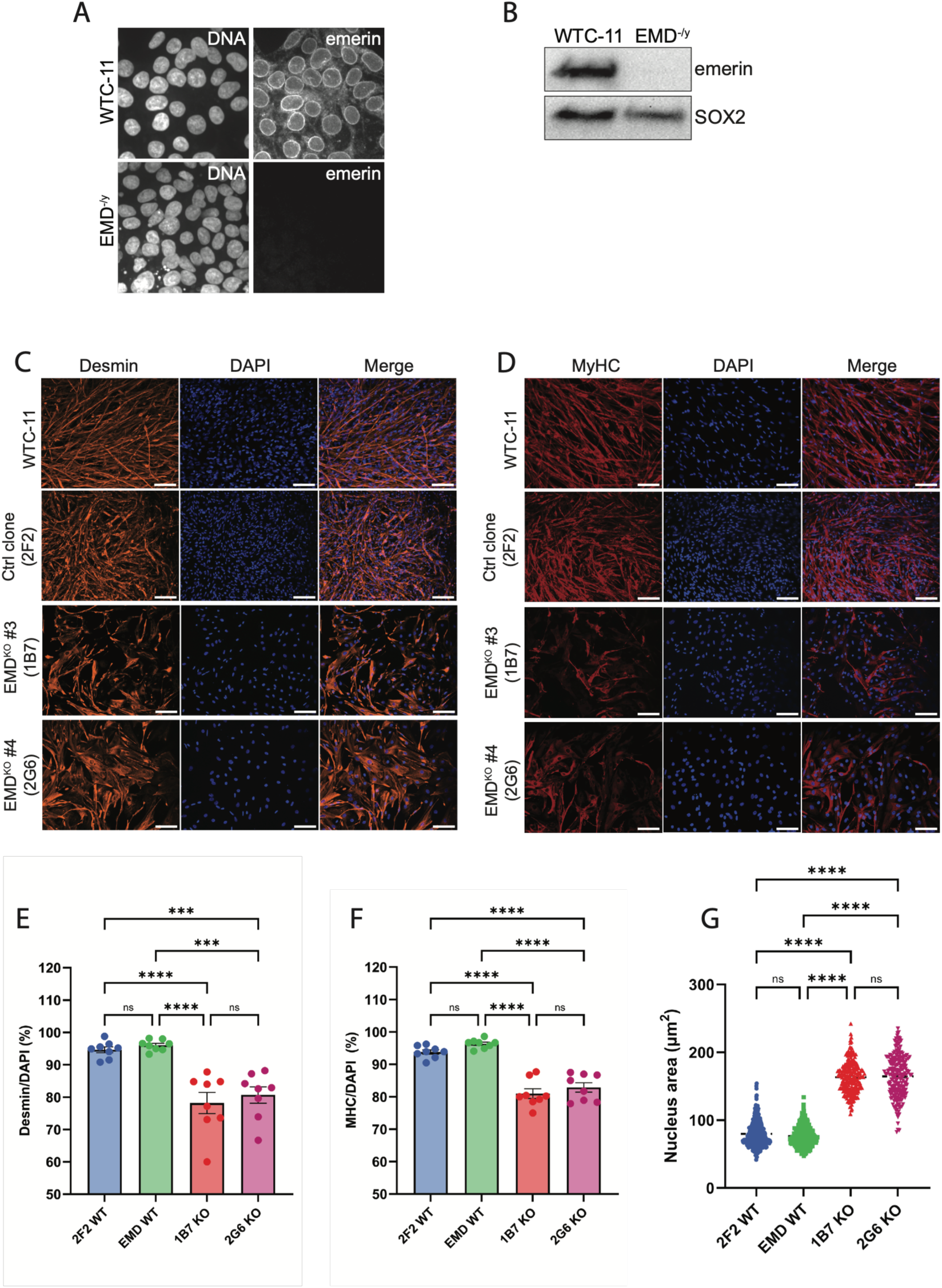
Additional EMD KO clones have a skeletal muscle differentiation defect. A-B) Representative EMD-/y iPSC clone lacking detectable emerin expression by immunostaining (A) and Western blotting (B). Note preserved expression of pluripotency marker SOX2. C-D) Representative images of Desmin-positive (C) and MyHC-positive (D) skeletal muscle myotubes in WTC-11 parental hiPSCs, CRISPR control (2F2), and two *EMD* KO clones (1B7 & 2G6). E-F) Skeletal muscle differentiation efficiency scored as % of cells positive for Desmin (E) and MyHC (F) for WTC-11, CRISPR control clone (2F2), and two *EMD* KO clones (1B7 & 2G6). G) Nucleus area calculated for each nucleus within each field of view across replicates in WTC-11, CRISPR control (2F2), and two *EMD* KO clones (1B7 & 2G6). Scale bar = 100 μm. Ordinary One-way ANOVA with multiple comparisons. ***p<0.001. ****p<0.0001.

**Supplementary Figure 5.**
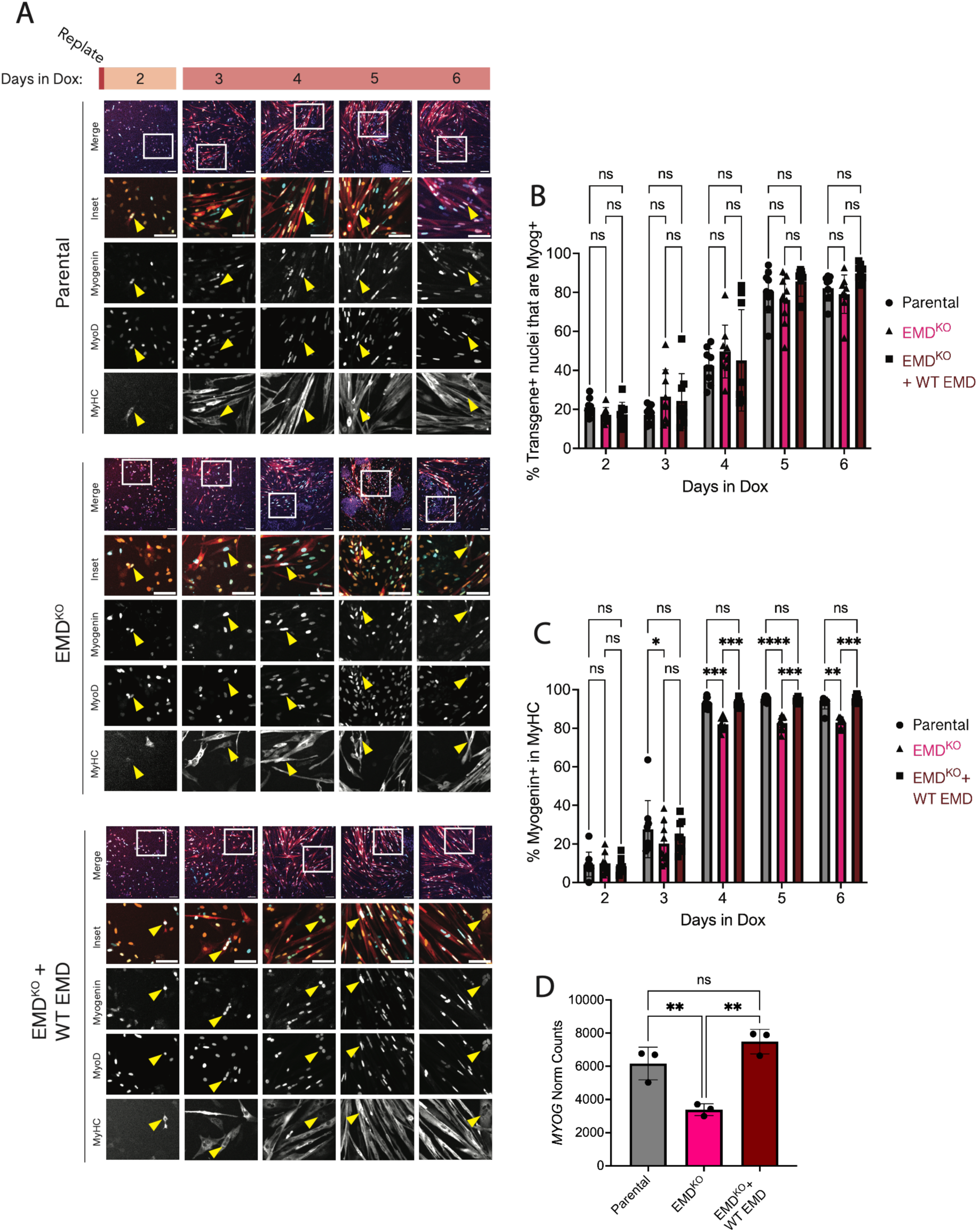
Myogenin expression is decreased in EMD KO iSMs. A) Time course of iSM differentiation, co-stained for myogenin, MyoD, and MyHC. Yellow arrowheads indicate transgene+/Myog+ population, quantified in (B). Quantification of transgene-positive nuclei that are also myogenin-positive (n = 3 biological replicates per genotype). C) Quantification of myogenin-positive nuclei within MyHC-positive myotubes in (A) (n = 3 biological replicates per genotype). D) Normalized transcript counts of *MYOG*. All scale bars = 100um. Proportions significantly different by two-way ANOVA in (B) and (C), one-way ANOVA in (D): **p < 0.01, *** p < 0.001, ****p < 0.0001.

**Supplementary Figure 6.**
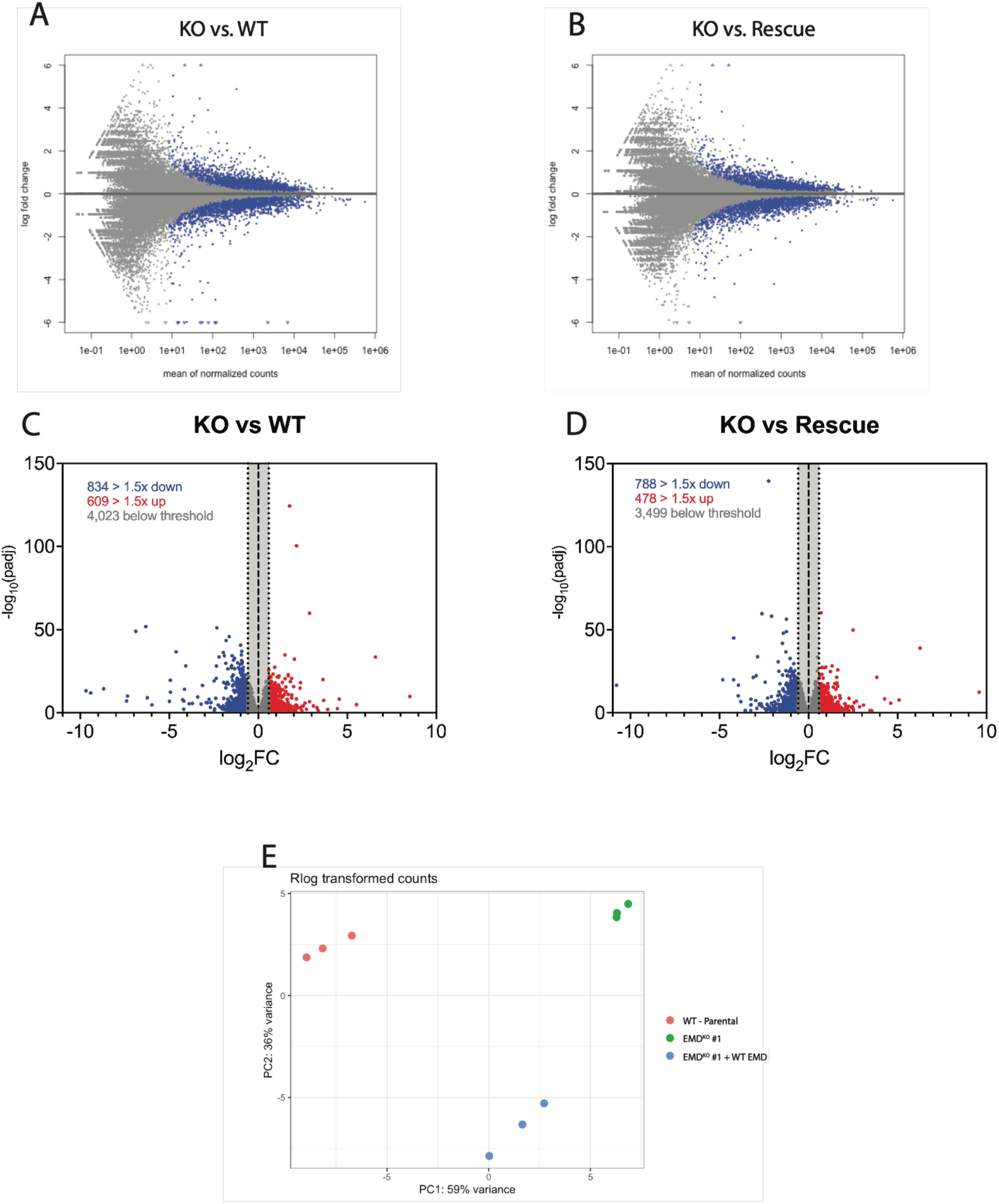
Analysis of transcriptional changes after EMD KO in Day 6 iSMs. A-B) MA plots comparing gene expression in KO vs. WT (A) and KO vs. Rescue (B). 5,466 differentially expressed genes in (A) and 4,765 in (B) with p-adj < 0.05 shown in blue. C-D) Log_2_FC of differentially expressed genes shown in (A) and (B), plotted against -log(p-adj). 609 genes are upregulated > 1.5-fold, 834 downregulated > 1.5-fold in (C). 478 genes are upregulated > 1.5-fold, 788 downregulated > 1.5-fold in (D). E) Principal component analysis (PCA) plot for three biological replicates across the three genotypes analyzed by RNA-seq.

**Supplementary Figure 7.**
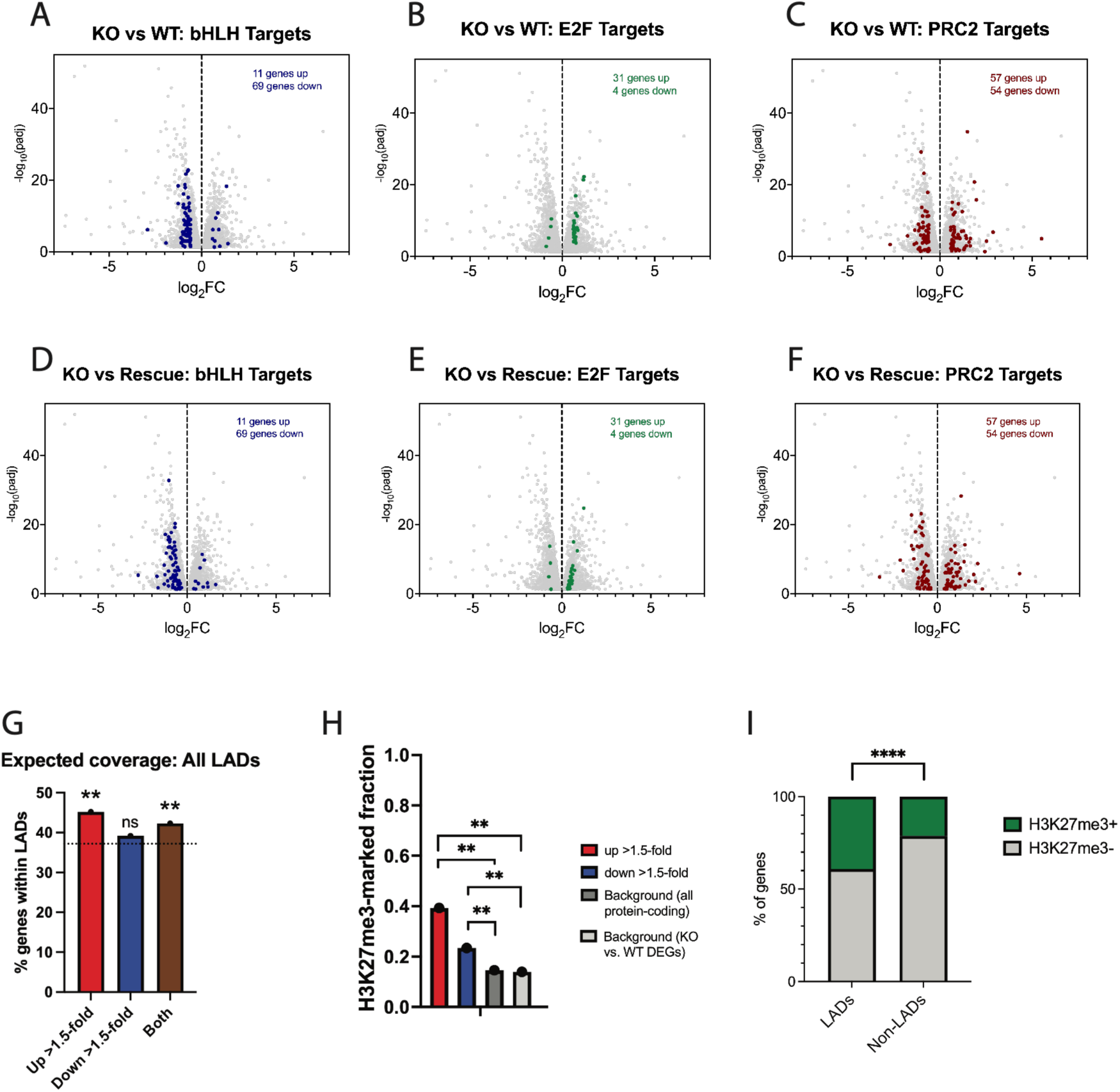
Analysis of rescue-sensitive gene distribution among transcription factor targets, LADs, and H3K27me3-positive regions. A-F) Log2FC of differentially expressed genes between KO vs. WT (A-C) and KO vs. Rescue (D-F), plotted against -log(p-adj). Gene targets of bHLH TFs (A & D), E2F TFs (B & E), and the PRC2 complex (C-F) are highlighted. G) LAD enrichment among up and downregulated rescue-sensitive genes, or the union of both. Dotted line represents background (average LAD coverage across all human protein-coding genes). H) Fraction of genes per set predicted to be H3K27me3-marked in human myotubes. Up and downregulated rescue-sensitive genes were compared to both an all protein-coding gene background and the full DEG list of KO vs. WT background. I) Percentage of all genes within LADs and non-LADs that are predicted to be H3K27me3-marked in human myotubes. All percentages significantly different when compared to background(s) by two-sided Fisher’s exact test: *p < 0.05, **p < 0.01, ***p < 0.001, ****p < 0.0001.

**Supplementary Table 1.**
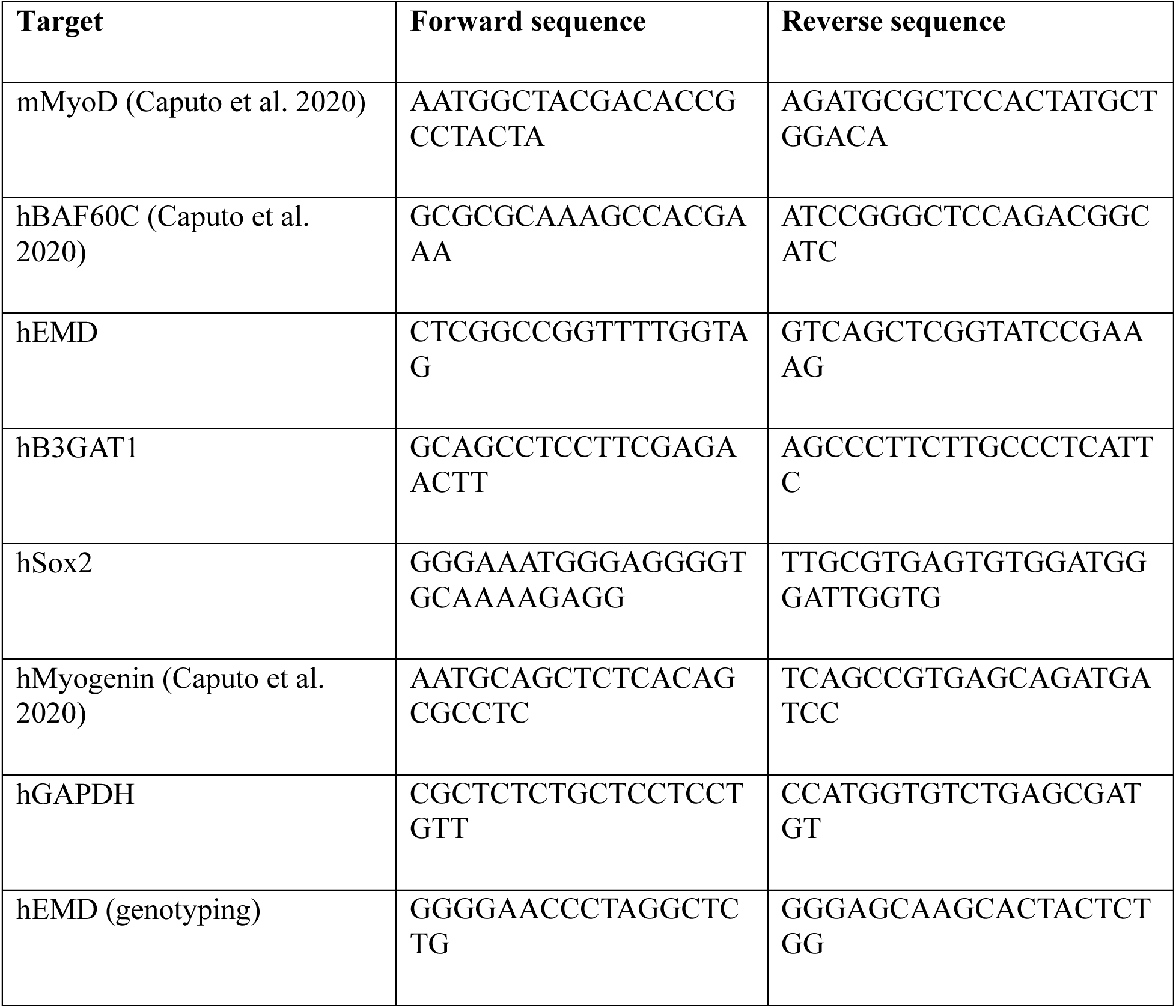
List of RT-qPCR and genotyping primers.

**Supplementary Table 2.** List of antibodies.

| Target | Isotype | Application | Dilution | Catalog No. |
| --- | --- | --- | --- | --- |
| MyoD | Mouse IgG1 | IF | 1:150 | BD 554130 |
|  |  | IP | 1:120 |  |
|  |  | WB | 1:1000 |  |
| MyoD | Rabbit | IF | 1:400 | Proteintech 18943-1-AP |
|  |  | WB | 1:2000 |  |
| BAF60C | Rabbit | IF | 1:100 | Proteintech12838-1-AP |
|  |  | WB | 1:1000 |  |
| MyHC | Mouse IgG2b | IF | 1:50 | DSHB MF 20 |
|  |  | WB | 1:100 |  |
| EMD | Rabbit | IF | 1:300 | Proteintech 10351-1-AP |
|  |  | WB | 1:2000 |  |
| p21 | Rabbit | IF | 1:400 | Cell Signaling<br>Technology 2947T |
|  |  | WB | 1:1000 |  |
| Myogenin | Mouse IgG1 | IF | 1:50 | DSHB F5D |
| Vinculin | Mouse IgG1 | WB | 1:4000 | Sigma-Aldrich V9264-100UL |
| LMNA | Mouse IgG2a | IF | 1:500 | Active Motif 39288 |
|  |  | WB | 1:1000 |  |
| Histone H4 | Rabbit | WB | 1:1000 | Active Motif 39070 |
| Pan-acetyl-K | Rabbit | WB | 1:1000 | Cell Signaling<br>Technology 9814S |
| E2A | Rabbit | WB | 1:1000 | Cell Signaling<br>Technology 4865S |
| Desmin | Mouse IgG1 | IF | 1:100 | Dako/Agilent M0760 |

